# ScreenDMT reveals linoleic acid diols replicably associate with BMI and stimulate adipocyte calcium fluxes

**DOI:** 10.1101/2023.07.12.548737

**Authors:** Jonathan M. Dreyfuss, Vera Djordjilovic, Hui Pan, Valerie Bussberg, Allison M. MacDonald, Niven R. Narain, Michael A. Kiebish, Matthias Blüher, Yu-Hua Tseng, Matthew D. Lynes

## Abstract

Activating brown adipose tissue (BAT) improves systemic metabolism, making it a promising target for metabolic syndrome. BAT is activated by 12,13-dihydroxy-9Z-octadecenoic acid (12,13-diHOME), which we previously identified to be inversely associated with BMI and which directly improves metabolism in multiple tissues. Here we profile plasma lipidomics from a cohort of 83 people and test which lipids’ association with BMI replicates in a concordant direction using our novel tool ScreenDMT, whose power and validity we demonstrate via mathematical proofs and simulations. We find that the linoleic acid diols 12,13-diHOME and 9,10-diHOME both replicably inversely associate with BMI and mechanistically activate calcium fluxes in mouse brown and white adipocytes in vitro, which implicates this pathway and 9,10-diHOME as candidate therapeutic targets. ScreenDMT can be applied to test directional mediation, directional replication, and qualitative interactions, such as identifying biomarkers whose association is shared (replication) or opposite (qualitative interaction) across diverse populations.

## Introduction

The incidence of pre-diabetes, metabolic syndrome, and type 2 diabetes are increasing word-wide and pose a critical need for improved therapeutics. A promising approach is activating brown adipose tissue (BAT), which increases the rate of systemic metabolism by oxidizing metabolic fuels like glucose and fatty acids and secreting signaling molecules. We and others have discovered that BAT can produce omega (ω)-6 and ω-3 polyunsaturated fatty acids (PUFAs)-derived bioactive lipids (i.e. lipokines) such as 12,13-dihydroxy-octadecaenoic acid (12,13-diHOME), which increases BAT activity (Leiria, Wang et al. 2019, Shamsi, Wang et al. 2021), skeletal muscle fatty acid uptake (Stanford, Lynes et al. 2018), cardiac function (Pinckard, Shettigar et al. 2021), and endothelial function to reduce atherosclerosis (Park, Li et al. 2022). Furthermore, 12,13-diHOME in mother’s milk is associated with lower infant fat mass (Wolfs, Lynes et al. 2021) and an open-label clinical trial has found that sustained BAT activation over 4 weeks improved systemic metabolism (O’Mara, Johnson et al. 2020).

We had previously identified 12,13-diHOME due to its inverse association with BMI (body mass index) in a cohort of 55 individuals (Study 1) with a broad range of BMI (Lynes, Leiria et al. 2017) and then showed that 12,13-diHOME mechanistically induces translocation of fatty acid transport proteins to the surface of cells (Lynes, Leiria et al. 2017). Other groups also observed a relationship between human obesity and 12,13-diHOME; however, the signaling pathway that drives this effect is unknown. To confirm this association in another study using analogous methods and gain the power to identify additional signaling lipids that benefit systemic metabolism, we recruited a new cohort.

In this work, we profile signaling lipids in a cohort of 83 individuals (Study 2) that was partitioned based on BMI into obese (BMI > 40 kg/m^2^) and non-obese (BMI< 30 kg/m^2^). We first test if we can replicate the significant, negative rank correlation between plasma concentration of 12,13-diHOME at room temperature and BMI in Study 2. Secondly, we examine if other lipids’ association with BMI replicates in both studies (Lynes, Leiria et al. 2017). Replication of two studies tests a mathematically equivalent null hypothesis as mediation analysis, because both settings evaluate two independent hypotheses (mediation tests if each lipid or analyte is both associated with the exposure, and separately if it is associated with the outcome given the exposure). Replication and mediation can be tested by calculating each analyte’s p-value as the analyte’s maximum p-value over the two studies (i.e. the “joint significance” or “MaxP” method*’s* p-value) and then adjusting these maximum p-values for multiple testing of many analytes (Benjamini, Heller et al. 2009). The MaxP p-values were found via simulations to control the error rate and have more power than many competing approaches (MacKinnon, Lockwood et al. 2002, Barfield, Shen et al. 2017).

However, such application of the MaxP method has two limitations: it does not account for direction, whereas it is natural to require an effect to have the same direction in both studies to be considered replicated, and there are improved methods of adjusting p-values in this context. We previously accounted for directionality in High-throughput mediation analysis (Hitman), which proposed a Directional MaxP Test (DMT) and use of empirical Bayesian linear regression modeling to improve power (Dreyfuss, Yuchi et al. 2021), although here we apply non-parametric methods to mirror our previous analysis of 12,13-diHOME (Lynes, Leiria et al. 2017). The directional MaxP test assessed mediation in a direction consistent with the exposure’s effect on the outcome and its p-value is half the (non-directional) MaxP p-value if the direction is consistent, whereas otherwise its p-value is one. DMT was mathematically proven to control the error rate and was found to have stronger p-values than the MaxP and other methods (Dreyfuss, Yuchi et al. 2021).

We improved MaxP’s adjusted p-values by developing ScreenMin (Djordjilović, Page et al. 2019), which reduces the multiple testing burden by filtering out analytes whose minimum p-value is so weak that their MaxP p-value could never be significant. This idea was further developed in AdaFilter (Wang, Gui et al. 2022), which also proposes a method to account for direction. There are other methods that improve upon MaxP adjusted p-values and can account for direction, such as radjust-sym (Bogomolov and Heller 2018), where analytes with strong p-values from each study are selected and only analytes selected from both studies are tested for replicability, and the empirical Bayesian method RepFdr (Heller and Yekutieli 2014).

## Results

### Development and validation of ScreenDMT

To identify which lipid’s association with BMI is replicated, we sought a test that accounted for direction and had the strongest adjusted p-values in terms of the false discovery rate (FDR). Because we cared about directionality it was natural to consider DMT. By comparison, it was recently shown that the MaxP test is the likelihood ratio test (LRT) for the non-directional mediation null hypothesis (Liu, Shen et al. 2022), which is mathematically equivalent to testing replication without regard for direction in two studies. The LRT has many beneficial properties such as often being optimal, which is why LRTs so often used in practice, e.g. the *t*-test for means. For directional mediation and replication, we mathematically prove that DMT is the LRT in Text S1. To our knowledge, analyses of replication and mediation had not been previously connected to analysis of qualitative interactions, which occurs when an effect changes sign between groups of subjects, e.g. a treatment benefits one group of patients while it harms another. Surprisingly, we find that directional replication and mediation are connected to qualitative interaction, because applying the DMT in search of effects that have opposite signs between groups is the LRT for qualitative interactions (Gail and Simon 1985, Hudson and Shojaie 2020).

We considered how to improve the adjusted p-values from our directional MaxP test with screening or filtering. The filtering approach in AdaFilter when applied to two studies uses the minimum p-value (p_**min**_) per analyte as the filtering p-value and the maximum p-value as the selection p-value. We cannot directly use AdaFilter by naively setting the DMT p-value (p_**DMT**_) as the selection p-value, since the selection p-value must exceed the filtering p-value, which would not always be true. Instead, we developed a new way to apply screening to our directional MaxP test, which we term “ScreenDMT.” ScreenDMT defines its selection p-value per analyte as the maximum of p_DMT_ and p_min_. We mathematically prove ScreenDMT to be valid for controlling the FDR and the family-wise error rate (FWER; the rate controlled by the Bonferonni method) in Text S1 for large sample sizes. For small sample sizes, the screening-based multiple testing adjustment is very slightly too strong, but this is where the DMT p-value, like similar statistical tests, is weak anyway. Like in AdaFilter (Wang, Gui et al. 2022), theoretical control of the FWER requires independence of p-values within each study, while control of the FDR for a large number of analytes allows weak dependence between analytes. Such weak dependence typically holds, for example, between gene loci and expression (Wang, Gui et al. 2022). Importantly, not all FDR methods theoretically allow dependence, such as RepFdr (Heller and Yekutieli 2014).

We compare ScreenDMT’s FDR and power via simulation against competing directional replication methods, including the directional approaches of AdaFilter (Wang, Gui et al. 2022), radjust-sym (Bogomolov and Heller 2018), RepFdr (Heller and Yekutieli 2014), and DMT (Figure 1). We also compare against a directional alternative to the MaxP test that was adapted from a meta-analysis due to Pearson (Owen 2009, Wang, Gui et al. 2022), which applies MaxP for each analyte to right-sided p-values of both studies, then applies MaxP to left-sided p-values, and doubles the minimum from the two applications. This procedure also underlies AdaFilter’s directional approach.

**Figure 1.**
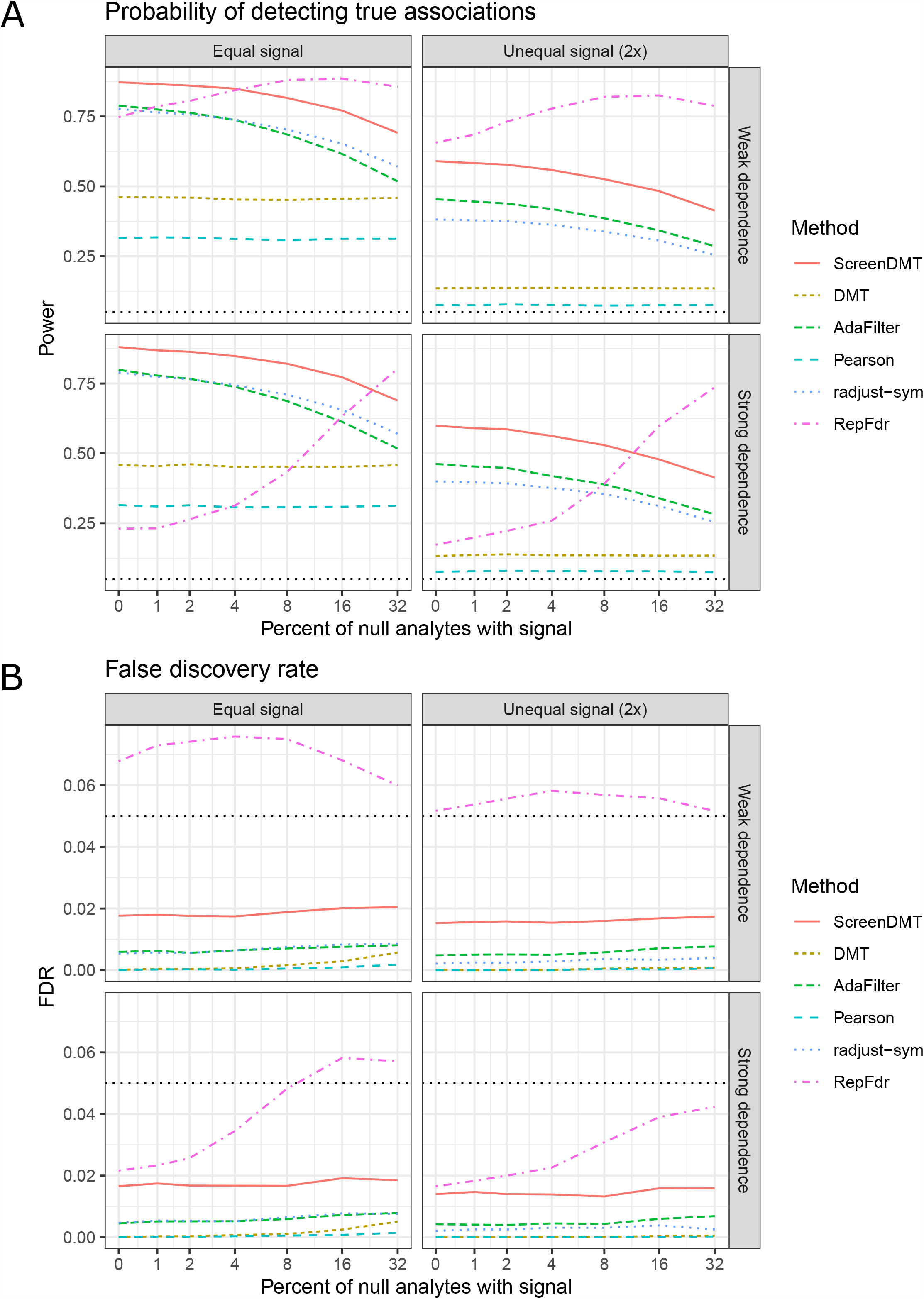
Comparison of ScreenDMT’s power and false discovery rate to competing approaches in replication of two datasets via simulation. Simulation of two datasets with strong or weak dependence between analytes when the signal strength or effect size per analyte is equal between the two datasets and when it is unequal. The x-axis represents the percent of analytes that do not truly directionally replicate but show some effect in one of the studies or show effect in both studies of opposite direction. (A) Probability of detecting true associations (power). (B) False discovery rate. The FDR threshold is 5%.

We see in Figure 1 that RepFdr often does not control its FDR below the nominal level, and apart from RepFdr, ScreenDMT retains the most power across the scenarios. We also see that DMT has more power than the approach adapted from Pearson (Owen 2009, Wang, Gui et al. 2022), which helps explain ScreenDMT’s power advantage over AdaFilter. We perform the analogous simulation for FWER using methods that account for directionality and provide FWER in Figure S1. We see that all methods properly control FWER, DMT is slightly more powerful than the method adapted from Pearson (Owen 2009, Wang, Gui et al. 2022), and ScreenDMT’s power is very similar but slightly stronger than AdaFilter’s.

### Inverse association of 12,13-diHOME with BMI replicates

We wanted to test if 12,13-diHOME negatively associates with BMI in the Study 2 cohort of 83 individuals (Table 1). Our previous association of 12,13-diHOME to BMI in Study 1 was assessed using Spearman rank correlation, so we also use a nonparametric test in Study 2 (Figure 2A, 2B). Since Study 2 is split into obese and non-obese subjects, we evaluate whether lipids are differentially abundant between groups with a non-parametric t-test (i.e.Wilcoxon rank sum or Mann-Whitney U test) (Figure 2C, 2D). We apply this test in a one-sided manner to test if 12,13-diHOME is lower in obese subjects. Since 12,13-diHOME had already been prioritized, its p-value does not need to be adjusted for multiple testing. We obtain a one-sided p-value of 0.00093 against a significance p-value threshold of 0.05, which demonstrates replication (Figure 2D).

**Table 1.** Anthropometrics of Study 2 cohort. Data are shown as mean for each group with the interquartile range in parentheses.

| Study group | Lean | Obese |
| --- | --- | --- |
| Age (years) | 52.1 (32.5-64.6) | 41.6 (31.3-51.9) |
| BMI (kg/m<sup>2</sup>) | 26.2 (24-28.8) | 45.1 (41.8-49.7) |
| Triglycerides (mmol/l) | 0.8 (0.6-03.5) | 5.3 (0.6-6.6) |
| Height (m) | 1.7 (1.6-1.8) | 1.7 (1.7-1.8) |
| Body weight (kg) | 75.5 (67.9-83) | 135.6 (117.9-150) |
| BMI (kg/m<sup>2</sup>) | 26.2 (24-28.8) | 45.1 (41.8-49.7) |
| Body fat (%) | 30.9 (23.1-38.6) | 46 (41.3-48.9) |
| Hemoglobin (mmol/l) | 8.3 (7.9-8.9) | 8.3 (7.9-8.9) |
| Erythrocytes (exp12/l) | 4.6 (4.3-4.8) | 4.7 (4.5-4.9) |
| Thrombocytes (exp9/l) | 259 (230.3-294) | 285.5 (244.5-320.3) |
| Leukocytes (exp9/l) | 6.6 (5.5-8) | 7.7 (6.4-8.8) |
| ALAT (μkat/l) | 0.4 (0.3-0.5) | 0.5 (0.4-0.6) |
| ASAT (μkat/l) | 0.5 (0.4-0.5) | 0.5 (0.4-0.6) |
| gGT (μkat/l) | 0.7 (0.3-0.7) | 0.7 (0.3-1.1) |
| Fasting plasma glucose (mmol/l) | 5.3 (4.8-5.9) | 5.8 (5.2-7.8) |
| HbA1c (%) | 5.6 (5.4-6) | 5.8 (5.3-6.6) |
| Fasting plasma insulin (pmol/l) | 31.3 (15.9-83.5) | 143.5 (79.2-185.4) |
| C-Peptid (nmol/l) | 1 (0.7-1.3) | 1.4 (0.9-2) |
| Creatinine (μmol/l) | 76 (59-85) | 67.5 (56.8-79.3) |
| Cholesterol total (mmol/l) | 5.5 (4.5-6.4) | 4.9 (4.3-5.3) |
| HDL-Cholesterol (mmol/l) | 1.3 (1.1-1.8) | 1.1 (0.9-1.5) |
| LDL-Cholesterol (mmol/l) | 3.1 (2.4-3.9) | 2.6 (2.1-2.9) |
| Total protein (g/l) | 72.3 (69.8-78) | 75.6 (73.2-84.1) |
| Albumin (g/l) | 45.9 (44.4-46.8) | 45.5 (43.6-46.7) |
| TSH (mU/l) | 1.5 (0.9-2) | 2.1 (1.4-3.3) |
| fT3 (pmol/l) | 4.8 (4.3-5.2) | 4.5 (4.3-5.3) |
| fT4 (pmol/l) | 15.7 (14-17.1) | 13.8 (13-15.6) |
| Cortisol (nmol/l) | 354 (212.9-549.7) | 396.3 (302.8-500.8) |
| Testosterone (nmol/l) | 1.1 (0.5-9) | 1.4 (0.9-5.5) |
| RRsys (mmHg) | 131 (124.3-139.3) | 143 (127-162) |
| RR dia (mmHg) | 84.5 (74.5-90) | 85 (80-94) |
| hsCRP (mg/dl) | 0.3 (0.2-1) | 2.2 (1.1-3.8) |

**Figure 2.**
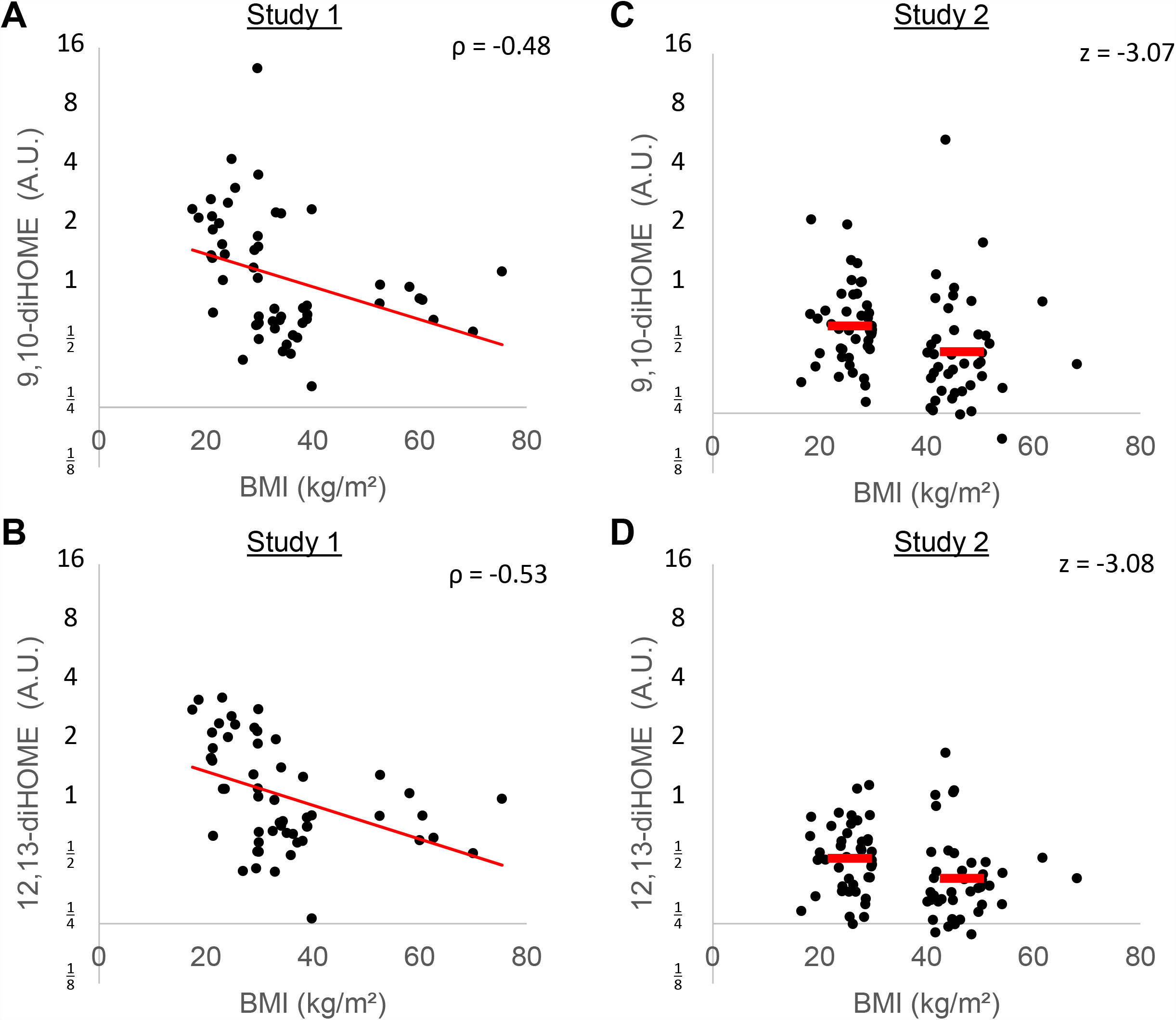
9,10-diHOME and 12,13-diHOME versus BMI in both cohorts. (A) Circulating 9,10-diHOME versus BMI in patients from the original cohort. (B) Circulating 12,13-diHOME versus BMI in patients from the original cohort. (C) Circulating 9,10-diHOME versus BMI in patients from the second validation cohort. (D) Circulating 12,13-diHOME versus BMI in patients from the second validation cohort. All lipid values were log_2_ transformed for graphing.

### Application of ScreenDMT to test replication of other lipids’ association with BMI

In addition to 12,13-diHOME, we measured a panel of 108 oxidized lipid species, including metabolites of linoleic acid, α-linolenic acid, arachidonic acid, dihomo-γ-linolenic acid, Docosahexaenoic acid and Eicosapentaenoic acid (Figure 3A). To assess if any lipids other than 12,13-diHOME have an association with BMI that replicates, we applied the non-parametric t-test to all the lipids except 12,13-diHOME and then assessed replication with ScreenDMT. The only other lipid that replicated was 9,10-dihydroxy-12Z-octadecenoic acid (9,10-diHOME), which is a regioisomer of 12,13-diHOME (Figure 3B), whose FDR is 5%. Whereas if we had tested replication with DMT, the FDR of 9,10-diHOME would be nearly double at 9%. We show the plots of 9,10-diHOME vs. BMI and of 9,10-diHOME vs. 12,13-diHOME in both cohorts in Figure 2.

**Figure 3.**
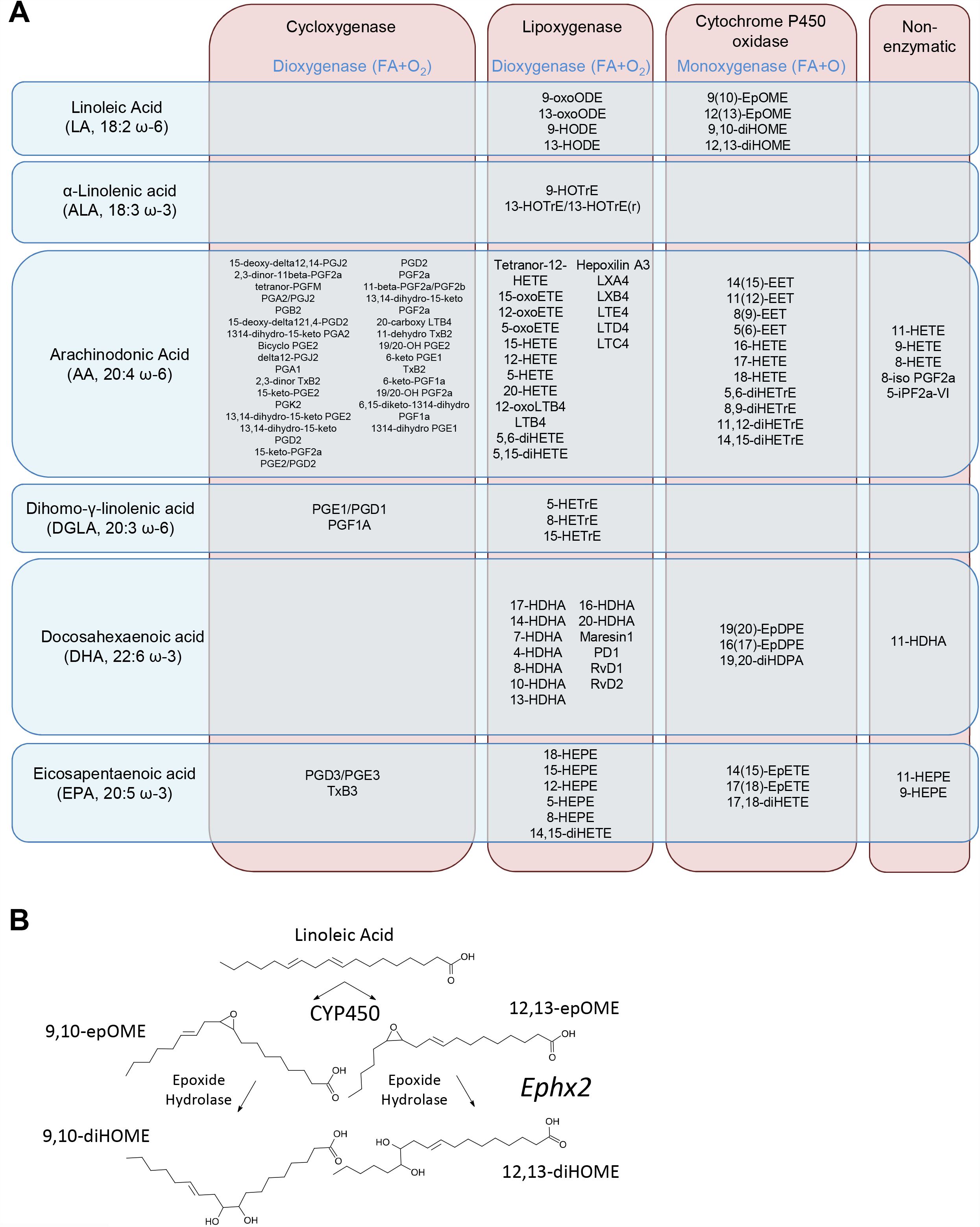
Biosynthesis of linoleic acid diols 9,10-diHOME. (A) Lipokines measured in our lipidomic panel shown as a product of their precursor fatty acids and the oxidative pathways that are the first step in their biosynthesis. (B) Biosynthesis of linoleic acid diols.

**Figure 4.**
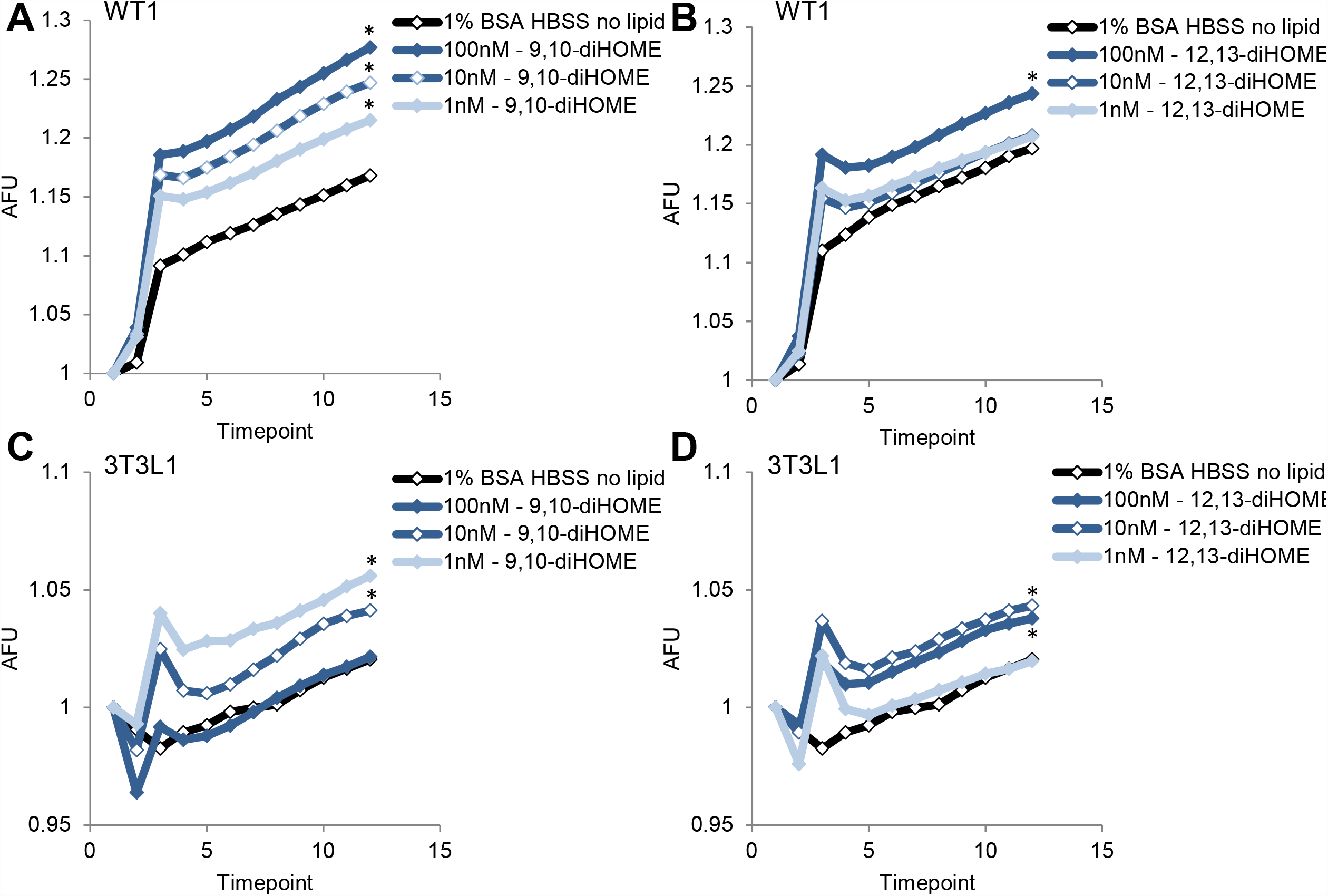
Linoleic acid diols activate calcium flux in cultured adipocytes. (A) FLUOFORTE quantification in differentiated WT1 brown adipocytes treated after timepoint 2 with different concentrations of 9,10-diHOME or vehicle control. (B) FLUOFORTE quantification in differentiated WT1 brown adipocytes treated after timepoint 2 with different concentrations of 12,13-diHOME or vehicle control. (C) FLUOFORTE quantification in differentiated 3T3-L1 white adipocytes treated after timepoint 2 with different concentrations of 9,10-diHOME or vehicle control. (D) FLUOFORTE quantification in differentiated 3T3-L1 white adipocytes treated after timepoint 2 with different concentrations of 12,13-diHOME or vehicle control. Each timepoint is approximately 50 seconds apart. Data are plotted as the normalized means ± s.e.m.; *n* = 6 wells per group; \**P* < 0.05 vs. vehicle by ANOVA with post-hoc Bonferroni test.

### 9,10-diHOME and 12,13-diHOME activate adipocyte calcium fluxes

12,13-diHOME and its stereoisomer 9,10-diHOME replicably associate with BMI, and we had previously shown that 12,13-diHOME increases cellular fatty acid uptake by inducing the translocation of fatty acid transporters FATP1 and CD36 to the cell surface; however, the signaling pathway that mediates this effect is unknown. 12,13-diHOME and 9,10-diHOME have both been found to activate ion flux in primary neurons and CHO cells (Green, Ruparel et al. 2016), so we measured calcium flux in adipocytes treated with each diol. We used in vitro differentiated WT1 mouse brown preadipocytes and in vitro differentiated 3T3-L1 mouse white preadipocytes to model brown and white adipocytes, respectively. Both cell lines were differentiated using a standard adipogenic induction media, and then loaded with Fura-2. After we recorded baseline calcium flux, cells were treated with Hanks Buffered Saline Solution (HBSS) containing 1% fatty acid-free bovine serum albumin (BSA) alone or containing different concentrations of linoleic acid diol, and calcium flux was monitored every 50 seconds for approximately 9 minutes in a plate reader (Figure 5). Both 9,10-diHOME and 12,13-diHOME triggered calcium influx in brown and white adipocytes, suggesting the signaling pathway activated downstream of these two lipids overlaps.

## Methods

### Study 2 Cohort

We included 84 individuals with (n = 41) or without (n = 42) obesity defined by a BMI > 30 kg/m^2^ (Table 1). Study participants were consecutively recruited to the Leipzig Obesity BioBank (LOBB). The study was approved by the local Ethics Committee of the University of Leipzig, Germany (Reg. numbers: 363–10-13122010 and 017–12-230112) and all subjects gave written witnessed informed consent before inclusion into the LOBB repository.

### Lipidomics

Plasma was analyzed by liquid chromatography tandem mass spectrometry (LC-MS/MS) to semi-quantitatively measure the concentrations of a panel of 113 signaling lipids. Plasma samples were thawed at room temperature and immediately placed on ice. Aliquots of 100 µL were taken and added to 300 µL of methanol (stored at -20 °C) for a protein crash. Ten µL of a mixture of 5 deuterated internal standards, each at 100 pg/µL, were spiked into the samples, then samples were vortexed for 10 seconds and stored at -20 °C overnight. Samples were then subjected to a solid phase extraction. C18 cartridges at 500 mg/6 mL (Biotage, Uppsala, Sweden) were conditioned with 10 mL of methanol followed by 10 mL of water. The samples were centrifuged at 14000 g, and the pH of the supernatant of the samples was adjusted by adding 3 mL of pH 3.5 water before loading the samples onto the C18 cartridges. The cartridges were washed with 5 mL water followed by 5 mL hexanes. Fractions were dried down under a stream of N2 gas, and reconstituted in 50 µL methanol:water (1:1, by vol). Samples were vortexed and then transferred to LC-MS vials for analysis.

Electrospray ionization (ESI) LC-MS/MS was performed on a QTRAP 6500 (Sciex, Framingham, MA, USA) coupled to an Agilent Infinity 1290 (Agilent, Santa Clara, CA, USA) LC system with a InfinityLab Poroshell 120 EC-C18 (4.6 × 100 mm, 2.7 µm; Agilent) analytical LC column with a column oven heated to 60°C. Ten µL of sample was injected at a flow rate of 400 µL/min and were separated by reversed phase chromatography with mobile phases A (100% H2O, 0.1% acetic acid) and B (100% MeOH, 0.1% acetic acid). The gradient started at 5% B and increased to 20% B by 3 minutes, then increased to 60% B by 8 minutes, increased to 90% B by 23 minutes, held at 90% B for 3 minutes, then returned to 5% B for the last 3 minutes. The total LC run time was 29 minutes. Samples were only acquired in negative polarity due to the chemical structure of the targeted lipids. The ESI source parameters were ion source gas 1 (GS1) 30, ion source gas 2 (GS2) 30, curtain gas (CUR) 30, ion spray voltage (IS) -4500, and temperature 500. The declustering potential (DP), entrance potential (EP), collision energy (CE), and exit potential (CXP) were tuned for each individual targeted lipid and internal standard. The MS method was a targeted scheduled MRM method. There were 101 MRM scans scheduled for optimized windows ranging from 120–360 seconds. The mass spectrometer acquisition time was set to 29 minutes. The LC-MS/MS data was acquired by Analyst 1.6.2 software (Sciex, Framingham, MA, USA) and processed with MultiQuant 3.0.1 (Sciex, Framingham, MA, USA) for peak integration. The targeted lipid species were measured semi-quantitatively and reported as area ratios of the peak area of the analyte divided by the peak area of the internal standard.

### Bioinformatics analysis of lipidomics data

We reanalyzed our previous lipidomics data from Study 1 (Lynes, Leiria et al. 2017) and Study 2’s lipidomics data using the R software. Both data sets included values of zero for missing data. Our only processing step was to filter out lipids with 20% or more missing values, which was applied to both datasets. To reproduce our previous study, we tested Spearman rank correlation with R function cor.test, which yielded the same coefficient and p-value as previously reported (Lynes, Leiria et al. 2017). For the Study 2 lipidomics dataset, we tested differential abundance between groups using the R function wilcox.test. Wilcoxon test z-scores were calculated from the wilcoxonZ function in the R package rcompanion. Replication FDRs from ScreenDMT were considered significant using FDR < 15%, as we used previously (Dreyfuss, Yuchi et al. 2021).

### Cell Culture

WT-1 mouse preadipocytes and 3T3-L1 mouse fibroblasts were cultured in high glucose DMEM supplemented with 10% FBS. When cells reached confluence, adipocyte differentiation was induced by using induction media (DMEM high glucose media with 10% FBS, 33 µM Biotin, 17 µM Pantothenate, 0.5 µM human insulin, 500 µM IBMX, 2 nM T3, 0.1 µM dexamethasone and 30 µM indomethacin) for 2 days, then cells were switched to a differentiation media (DMEM high glucose media with 10% FBS, 0.5 µM human insulin, 2 nMT3) for a final 7 days. After fully differentiated, adipocytes were washed with PBS and starved for 1 hour in DMEM.

### Calcium Flux Analysis

FLUOFORTE fluorescence was measured in adipocytes according to manufacturer’s directions (Enzo Scientific, Farmingdale, NY, USA). Briefly, starved adipocytes were incubated with FLUOFORTE Dye-loading solution for 1 hour at room temperature, then baseline fluorescence was read on a plate reader. After treatment with vehicle control or different concentrations of lipid diol, FLUOFORTE fluorescence was read every 50 seconds.

### Replication simulations

We assessed the performance of methods that provide adjusted p-values by simulating two studies with the same 1,000 features or analytes following the simulations in (Djordjilović, Hemerik et al. 2022). We simulated normally distributed statistics with variance of one per analyte per study. The statistics for analytes without signal were simulated with mean zero in both studies. In the Equal Signal simulation, the statistics for analytes that were truly directionally replicated were simulated with a mean of 3.5 or -3.5 in both studies, and these comprised 5% of the analytes. The statistics for analytes that were not directionally replicated but had some signal (“Percent of null analytes with signal”) were simulated with mean zero in one study and mean magnitude of 3.5 in the other, or else they were simulated with a mean of 3.5 in one study and -3.5 in the other study. In the Unequal Signal setting, the mean magnitudes for one study was 2.66 while the other study had mean magnitude of 4.66. Like the AdaFilter simulations (Wang, Gui et al. 2022), the statistics were simulated under weak dependence, where analytes were correlated with correlation coefficient of 0.5 to other features in their block with block size of 10, whereas under strong dependence, features were similarly correlated but in block sizes of 100. Features were randomly assorted to blocks.

For each setting 1000 simulations were run. RepFdr using its default parameters was too slow to run this many simulations, so the tolerance specified in RepFdr’s package vignette was used. So that RepFdr calculates FDRs when it estimates the fraction of nulls to be one in both studies, we modified it to calculate each analyte’s FDR as one instead of producing an error. For the FWER case, we do not compare to (Bogomolov and Heller 2018) because its replication implementation radjust-sym does not include a FWER procedure and its mediation implementation does not account for direction (Sampson, Boca et al. 2018).

## Discussion

We test replication which has been defined as obtaining the same result in new data (Nosek, Hardwicke et al. 2022). Although often used synonymously, “reproducibility” can be defined as obtaining the same result on the same data with the same analysis (Nosek, Hardwicke et al. 2022), whereas meta-analysis combines statistics from datasets, which can yield significance due to a strong effect in a single study. Replication is fundamental to science and its mathematical treatment has been found to have numerous applications (Benjamini, Heller et al. 2009). We focus on replication in a concordant direction and show that our test for directional replication is mathematically equivalent to testing for qualitative interactions and directional mediation.

An example of a qualitative interaction would be a pair of genes that are positively associated in healthy participants and negatively associated in those with disease. Finding such pairs is important for identifying the gene interactions causing disease and other questions in differential network biology (Ideker and Krogan 2012). Another example would be a biomarker being positively associated with an outcome in one population, such as those of European ancestry, while being negatively associated in an under-represented population.Although a limitation of our work is that Study 1 and Study 2 are comprised of people of European ancestry, our method can powerfully identify whether linoleic acid diols inverse associations to BMI (or any other findings in Europeans) are consistent or of the opposite direction across populations, promoting equity in medicine (Sirugo, Williams et al. 2019).

We demonstrated the value of mediation accounting for direction in a previous study where Hitman, which is based on DMT, identified growth hormone receptor as a mediator of gastric bypass surgery’s reduction of glycemia (Dreyfuss, Yuchi et al. 2021). We found that surgery reduced GHR levels, and lower GHR was associated with lower glycemia. Mediation methods that do not account for direction would have called the metabolite retinol as a mediator, even though surgery reduced retinol levels, and lower retinol was associated with glycemia being increased. Thus, surgery’s apparent action via retinol would be for glycemia to increase. This type of “inconsistent” mediation is unlikely and does not help explain mechanism of action, so our directional MaxP test saves power by assigning such an analyte a p-value of one. ScreenDMT improved upon the directional MaxP test with more efficient adjusted p-values (Wang, Gui et al. 2022). These improved adjusted p-values correspond to selection p-values that are based on, but not necessarily the same, as the directional MaxP test p-values. Thus, at times, the adjusted p-values may not have the same order as the DMT p-values. As usual when testing many hypotheses, significance should be based on adjusted p-values.

A possible reason that both 12,13-diHOME and 9,10-diHOME levels are associated with BMI is their shared biosynthetic pathways downstream of linoleic acid. Linoleic acid is a polyunsatured omega-6 fatty acid that is one of two essential fatty acids for humans and can be used as an energy source (Carneheim, Cannon et al. 1989). However, it can also be converted to arachinodonic acid and metabolized into a diverse group of signaling molecules, including the linoleic acid diols (Lynes, Kodani et al. 2019, Abe, Oguri et al. 2022). Linoleic acid diols are synthesized in a two-step process that begins with oxidation of linoleic acid at either the double bond at the 9,10 position or the 12,13 position, resulting in one of two regioisomer epoxides. This step is catalyzed by Cytochrome P450 oxidases, however little is known regarding the specific identity of the enzyme(s) that catalyze this step or if and how the position of the epoxide moiety is determined. Biosynthesis of 12,13-diHOME from its parent epoxide is mediated by an epoxide hydrolase, most likely soluble epoxide hydrolase encoded by the gene Ephx2, as mice lacking Ephx2 have circulating levels of 12,13-diHOME 8-fold lower than wildtype or Ephx1 knockout controls (Edin, Hamedani et al. 2018). Similarly, 9,10-diHOME is generated from an epoxide precursor called 9,10-epOME, however only genetic ablation of both Ephx2 and Ephx1, which encodes the microsomal isozyme of epoxide hydrolase, is sufficient to lower the levels of circulating 9,10-diHOME (Edin, Hamedani et al. 2018).

In mice and cells, 12,13-diHOME can activate changes in metabolism by recruiting fatty acid transport proteins to the cell surface, however the signaling pathway or pathways that mediate this effect are unknown. A candidate transport protein is CD36 and translocation of CD36 can be activated by calcium flux (Angin, Schwenk et al. 2014). Since both 12,13-diHOME and 9,10-diHOME can activate changes in membrane polarization (Green, Ruparel et al. 2016) and we have recently shown that 12,13-diHOME can activate calcium flux in bovine endothelial cells (Park, Li et al. 2022), we tested whether both lipids in their physiological range from 1-100nM (Lynes, Leiria et al. 2017, Vasan, Noordam et al. 2019, Kulterer, Niederstaetter et al. 2020) have a similar effect on adipocyte calcium flux and found that they do. The beneficial effect of 9,10-diHOME is historically surprising, because it was originally tested in the micromolar range where it was found to cause mitochondrial damage in leukocytes, so it was named Leukotoxin diol. However, given our new findings there is an evidence base to test the beneficial effects of 9,10-diHOME on systemic metabolism and given the overlap between signaling that we observed with 12,13-diHOME, the signaling pathway targeted by these two molecules could prove to be a valuable therapeutic target in the treatment of obesity and its sequalae.

## Figure Legends

**Figure S1.**
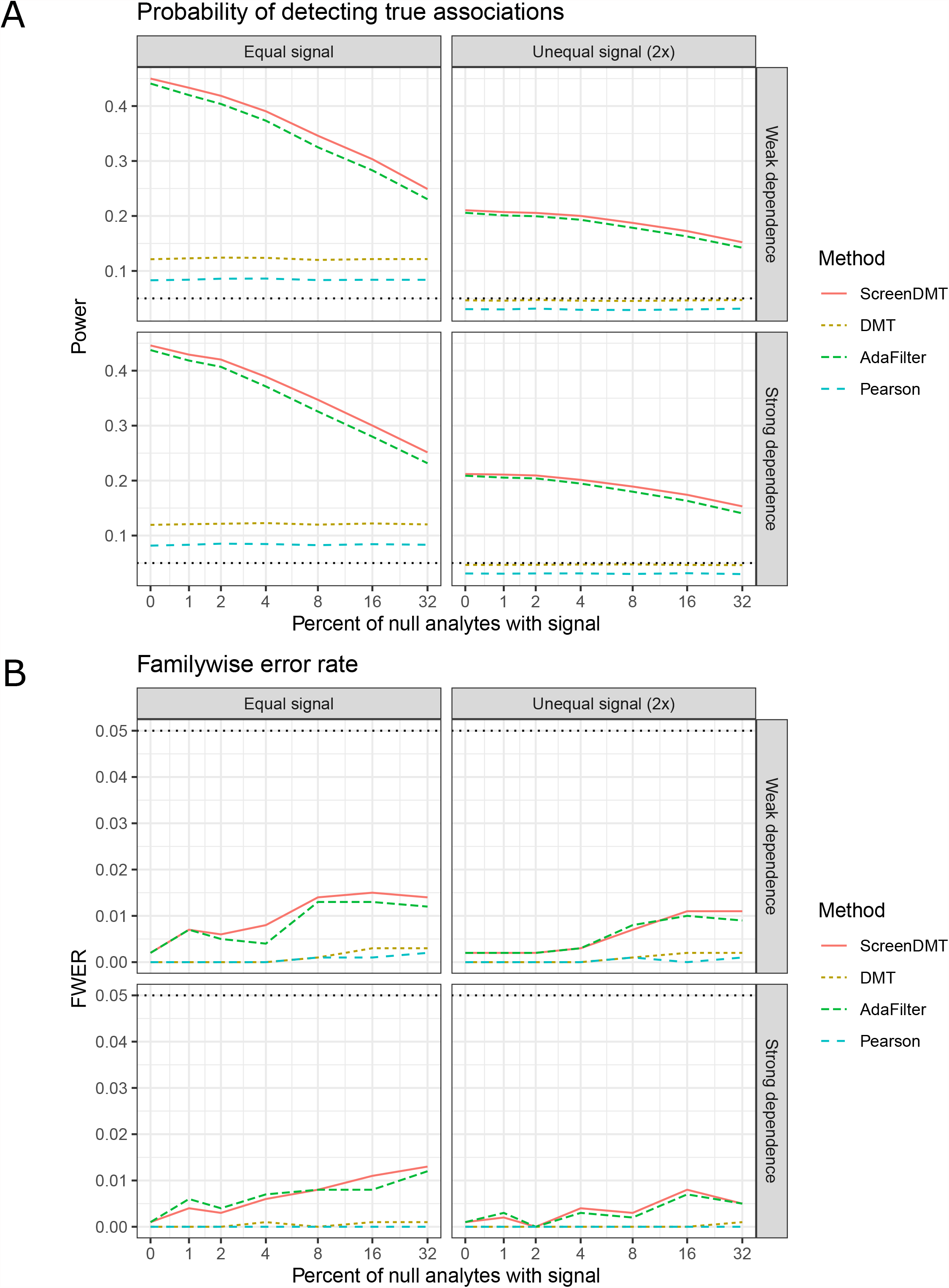
Comparison of ScreenDMT’s power and family-wise error rate to competing approaches in replication of two datasets via simulation. Simulation of two datasets with strong or weak dependence between analytes when the signal strength per analyte is equal between the two datasets and when it is unequal. The x-axis represents the percent of analytes that do not truly directionally replicate but show some effect in one of the studies or show effect in both studies of opposite direction. (A) Probability of detecting true associations (power). (B) Family-wise error rate (FWER). The FWER threshold is 5%.

## Author Contributions

Conceptualization: JMD, MDL; Methodology: JMD, VD, HP, MDL; Software: JMD, VD, HP; Validation: JMD, VD, MDL; Investigation: JMD, VD, VB, AMM, MDL; Writing – Original Draft: JMD, MDL. Writing – Review & Editing: VD, HP, Y-HT; Supervision: MAK, NRN, Y-HT.

## Data Availability Statement

Lipidomics measurements for both studies and sample metadata is freely available at https://github.com/jdreyf/screendmt-dihome-replication, which also contains R code to reproduce this paper’s lipidomics results and simulations. The code relies on the freely available R package DirectionalMaxPTest (https://github.com/jdreyf/DirectionalMaxPTest), which implements DMT and ScreenDMT. All datasets generated for this study are included in this article.

## Text S1: ScreenDMT procedure and validation

### Problem statement

Let *ω*_1_ and *ω*_2_ be two finite real valued parameters. In the mediation setting *ω*_1_ and *ω*_2_ represent the exposure-mediator and the mediator-outcome effect, while in the replication setting *ω*_1_ and *ω*_2_ represent parameters pertaining to the same biological entity measured in two studies. Whether we are interested in directional mediation or directional replication across two studies, without loss of generality we can consider the problem of detecting mediation in a direction consistent with the exposure increasing the outcome or replication in the same direction across two studies. Thus, we are interested in testing the directional null hypothesis:

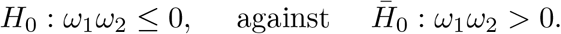

The null hypothesis can alternatively be written as:

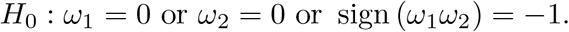

We assume that we have 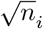-consistent and asymptotically normal estimates 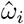 of *ω*_*i*_ for *i* = 1, 2 such that

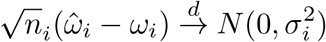

where the convergence is convergence in distribution and 0 *< σ*_*i*_ *<∞* for *i* = 1, 2, where *σ*_*i*_ is known. Then we can define component *p*-values *P*_1_ and *P*_2_ for testing *H*_1_ : *ω*_1_ = 0 and *H*_2_ : *ω*_2_ = 0, respectively,

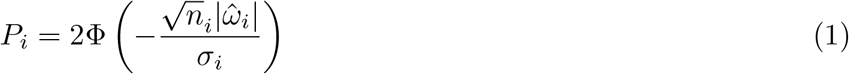

where Φ(*·*) is the standard normal cumulative distribution function. The procedures below are valid for large samples if consistent estimates of *σ*_*i*_ are used instead. We further assume that 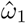 and 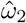 are independent (i.e. 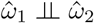), a plausible assumption in both the mediation and the replication setting.

To simplify notation we let

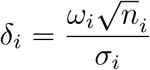

for *i* = 1, 2 so that Equation 1 becomes 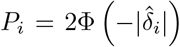. This is without loss of generality, since in any dataset 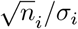 is a constant coefficient of 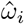. The constant is always positive, so *δ*_*i*_ could replace *ω*_*i*_ in the problem statement. We assume *ω*_*i*_ and *σ*_*i*_ are fixed, so |*δ*_*i*_|*→ ∞*implies *δ*_*i*_ ≠ 0 and *n*_*i*_*→ ∞*. This problem’s null (white) and alternative (grey) regions are shown in the figure below.

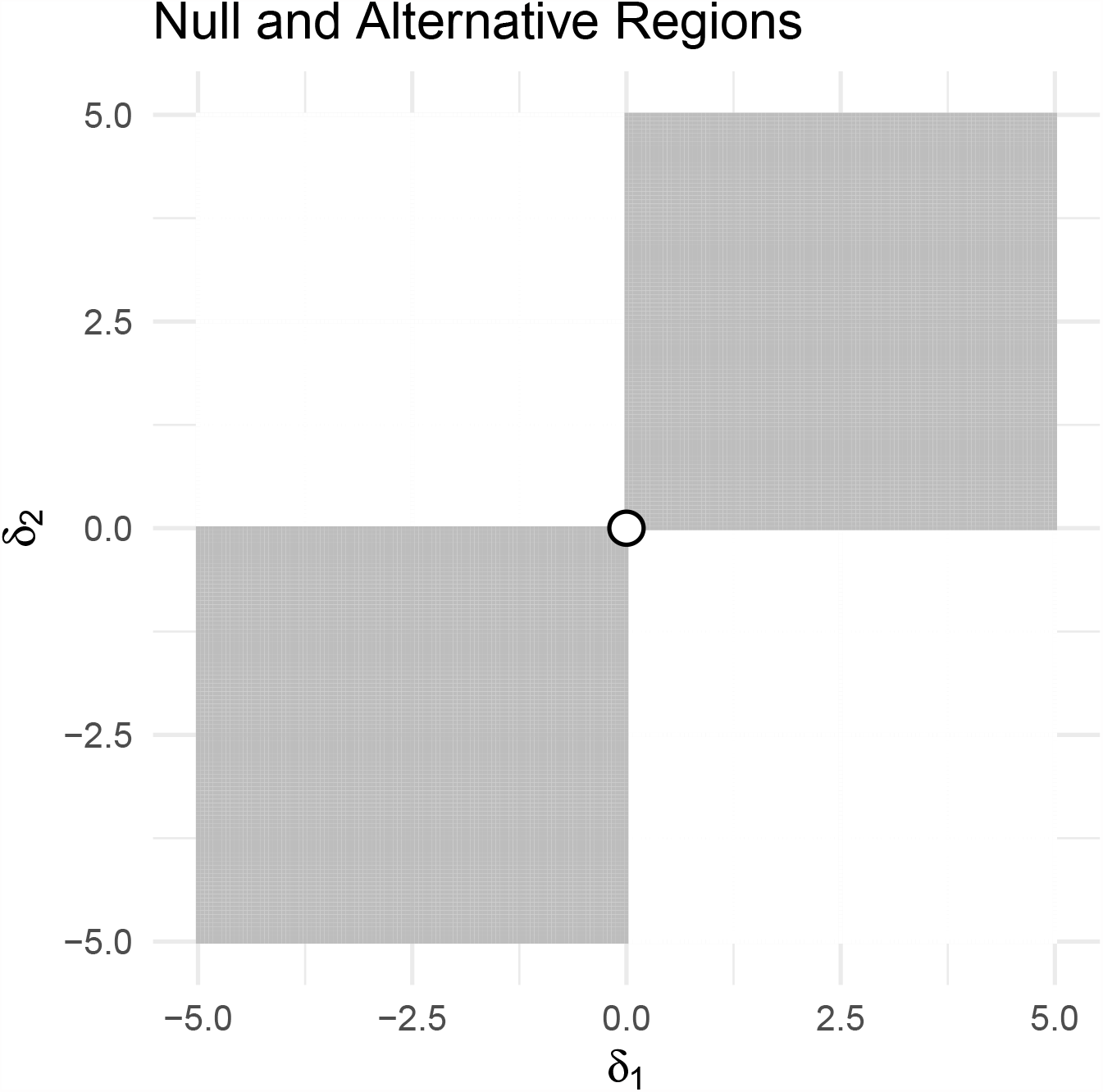

### Directional MaxP test

#### Previous result

The following result given in Dreyfuss et al. (2021) regarding the directional MaxP test introduced in the Hitman method specifies how a *p*-value for *H*_0_ can be obtained from both two-sided *p*-values associated to the two parameters under study.

Let *P*_1_ and *P*_2_ be the two valid component *p*-values for testing *H*_1_ : *ω*_1_ = 0 and *H*_2_ : *ω*_2_ = 0 and let 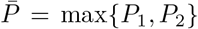 and 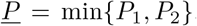. Further, let 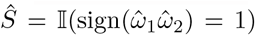 indicate if the signs of the estimates 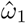 and 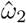 are the same, where 𝕀(*·*) is the indicator function which is one if its argument is true and zero otherwise. Then

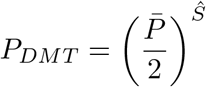

is a valid *p*-value for testing *H*_0_.

We showed this by demonstrating that ℙ(*P*_*DMT*_ *≤u*) *≤u* for 0 *≤u ≤*1. The case of *u* = 0 is trivial and we identified that under the null hypothesis 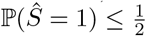 and then focused on the more challenging case of 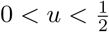. We also showed that 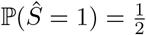 if and only (*δ*_1_, *δ*_2_) is on the boundary of the parameter space where *δ* _1_ = 0 ∪ *δ*_2_ = 0. Since *δ*_1_ and *δ*_2_ are d symmetrically, without loss of generality we considered the null region 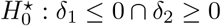. On the boundary of 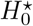, we found that ℙ(*P*_*DMT*_ ≤ *u*|*δ*_1_ = 0, *δ*_2_ = 0) = 2*u*^2^ *< u*, whereas if one parameter is zero and the magnitude of the other tends to infinity (i.e. the parameter is nonzero and the sample size tends to infinity), we saw that ℙ(*P*_*DMT*_ ≤ *u*)*→u*, which is the least favorable null. Gail and Simon (1985) came to the same conclusion, which they proved using Schur-convexity. We also found that when (*δ*_1_, *δ*_2_)*→* (*−∞, ∞*) then P(*Ŝ* = 1)*→*0 so ℙ(*P*_*DMT*_*≤u*)*→*0, which is the global minimum.

### Proposition 1: The directional MaxP test is the Likelihood Ratio Test (LRT) Proof

This problem is related to the problem of finding subsets of samples with effects of different signs (i.e. qualitative interactions, QI), whose LRT was shown by Gail and Simon (1985) and nicely presented in Hudson and Shojaie (2020). In that problem, 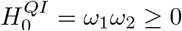 is tested against 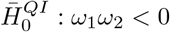, so the null region of the parameter space is 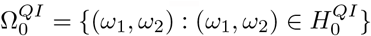. This is also the form of the problem in directional mediation when the exposure decreases the outcome (Dreyfuss et al. 2021).

For our problem, consider applying a LRT based on the asymptotic sampling distribution of 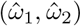. The LRT rejects *H*_0_ for large values of:

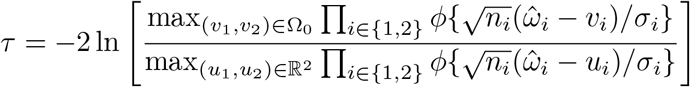

where Ω _0_ = {(*x*_1_, *x*_2_) : (*x*_1_, *x*_2_) *∈ H*_0_} is the null region of the parameter space (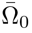 is the alternative region) and *ϕ*(·) is the standard normal density function. The denominator is maximized at 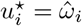 for both *i* = 1, 2. Then,

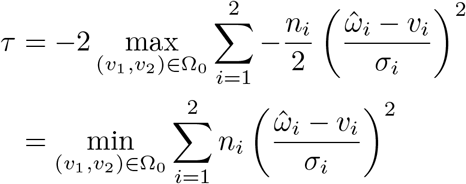

If 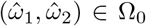, then we will have 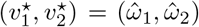, yielding *τ* = 0. So we next consider points under the alternative hypothesis. Without loss of generality, consider that 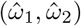 is in quadrant I, where both parameters are non-negative. Then the closest (*v*_1_, *v*_2_) can be to 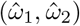 is to be on the boundary of quadrant I, where for one of *i* = 1 or *i* = 2 we have that 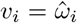 and for the other *i, v*_*i*_ = 0. The minimization of the sum will select the study with the smallest 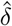:

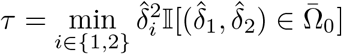

where the *p*-value of the LRT is computed as 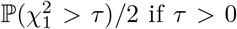, otherwise the *p*-value is one. This is equivalent to computing the *p*-value as

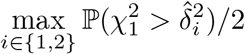

if 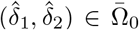 or otherwise one, which is asymptotically equivalent to the directional MaxP test, so the DMT is the LRT.

### ScreenDMT: multiple testing of directional hypotheses with screening

When considering multiple hypotheses (e.g. for many analytes, lipids, genes), we propose the following two-step multiple testing procedure. The ScreenDMT procedure includes an additional step based on AdaFilter (Wang et al. 2022) to calculate the adjusted p-values.

1. Apply the directional MaxP test per potential analyte to obtain *P*_*DMT*_.
2. Calculate adjusted p-values using AdaFilter with filtering p-value Λ = <u>*P*</u> and selection p-value Ξ = max{*P*_*DMT*_, <u>*P*</u>}. This can be used for calculating adjusted p-values that control the FDR or the FWER.

Dreyfuss et al. (2021) provided an option in step 1 to calculate valid p-values of these parameters in omics data using the R package Limma (Ritchie et al. 2015), which has been found to improve power across omics datasets. The same option is available for ScreenDMT.

The key property that AdaFilter relies on is a form of conditional validity (Wang et al. 2022). Assuming that for each analyte *P*_1_ is independent of *P*_2_, the property states that when *H*_0_ is true:

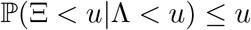

holds for any fixed u > 0 whenever ℙ(*P < u*) *>* 0.

### Proposition 2: ScreenDMT satisfies conditional validity in asymptotic sample sizes

**Proof**

To demonstrate conditional validity, we must show that this holds on the boundary of the parameter space where *ω*_1_ = 0 *∪ ω*_2_ = 0 and off the boundary where the first partial derivatives are zero. We have,

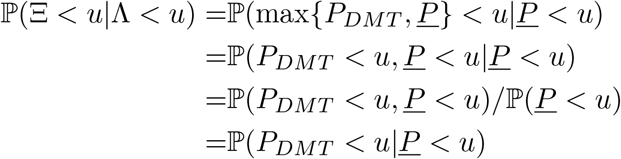

For *u≥*1, we trivially have ℙ (*P*_*DMT*_ *< u*|*<u>P</u> < u*) *≤u*. We know that *P*_*DMT*_ and *<u>P</u>* are symmetric in *ω*_1_ and *ω*_2_, so without loss of generality we only consider the null region 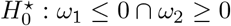. For 0 *< u <* 1, using the law of total probability, we continue

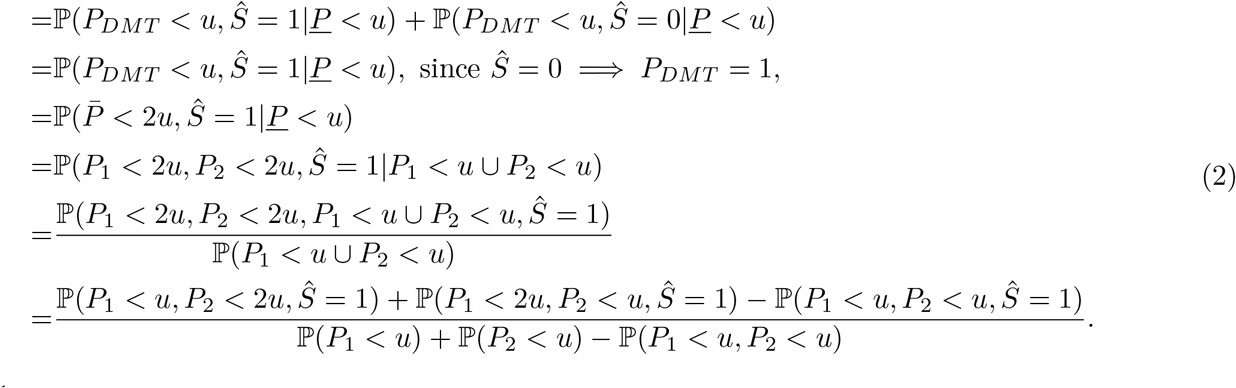

For 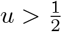we know that *P*_*i*_ *<* 2*u* for *i* = 1, 2, so Equation 2 becomes

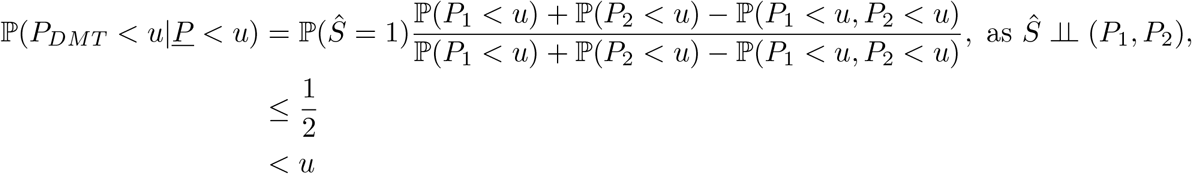

where 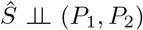 because *P*_*i*_ is calculated based on 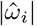, which is independent of *Ŝ*, and we showed the bound 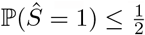previously (Dreyfuss et al. 2021).

We now consider ℙ(*P*_*DMT*_ *< u*|*<u>P</u> < u*) under 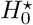for 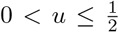both on the boundary of the parameterspace where *ω*_1_= 0 *∪ ω*_2_ = 0 and then over the whole parameter space.

#### Boundary of parameter space

Under 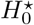for 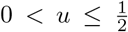when *ω*_1_ = 0 ∪ *ω*_2_ = 0 we know that 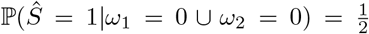 and ℙ(*P*_*i*_ *< u*|*ω*_*i*_ = 0) = *u*. Without loss of generality, let *ω*_2_ = 0 and *ω*_1_ *≤* 0, so ℙ(*P*_2_ *< u*|*ω*_2_ = 0) = *u* and ℙ(*P*_1_ *< u*|*ω*_1_ *≤* 0) = *F* (*u*) *≥ u*. For these parameter values, Equation 2 becomes

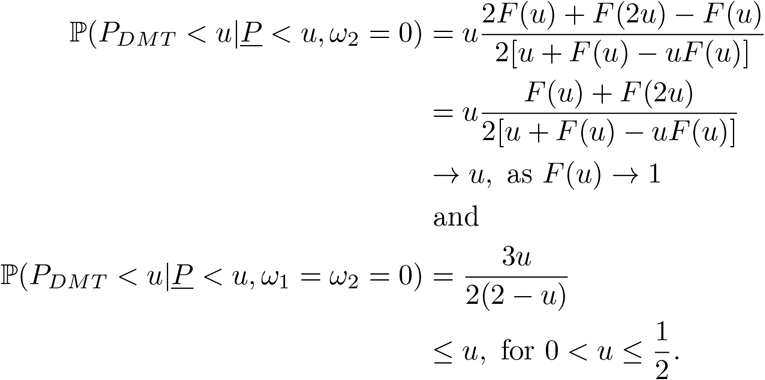

*F* (*u*) *→* 1 for *δ*_2_ = 0 as *δ*_1_ *→ −∞* as per Equation 1 which occurs as *n*_1_ *→ ∞*.

#### Whole parameter space

We continue Equation 2 for 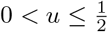using the definition of our component p-values from Equation 1,

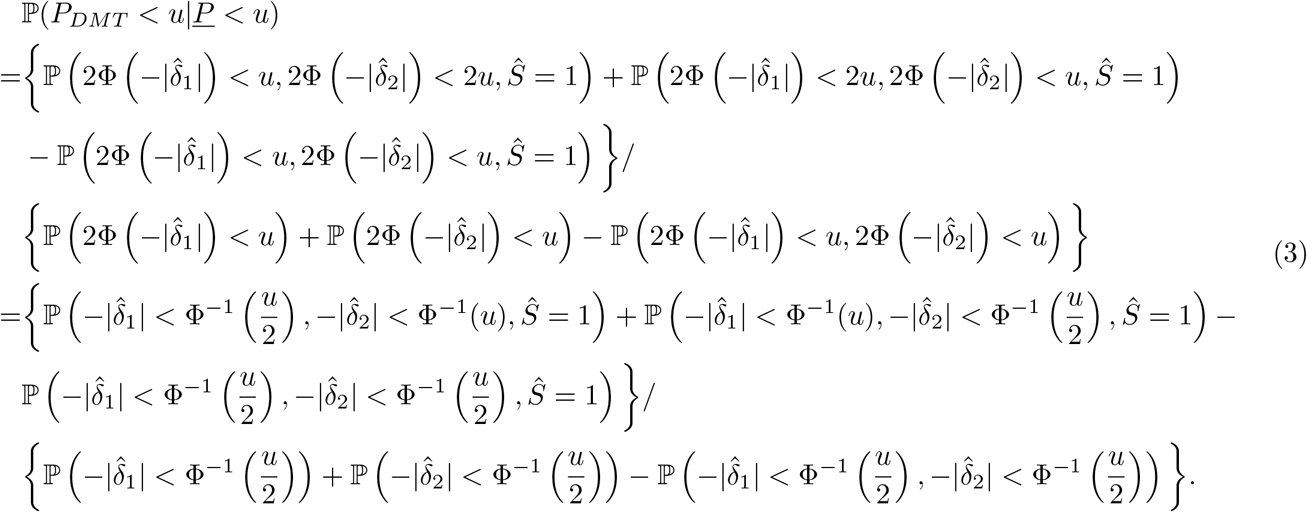

We calculate these probabilities according to the law of total probability by integrating over the realized values (*x*_1_, *x*_2_) of the random estimates 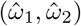. This involves the joint density

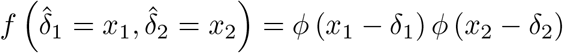

which follows from 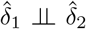, where *ϕ*(*·*) is the standard normal density. For example, for the first term in the numerator of Equation 3 with fixed *δ*_*i*_ for *i* = 1, 2,

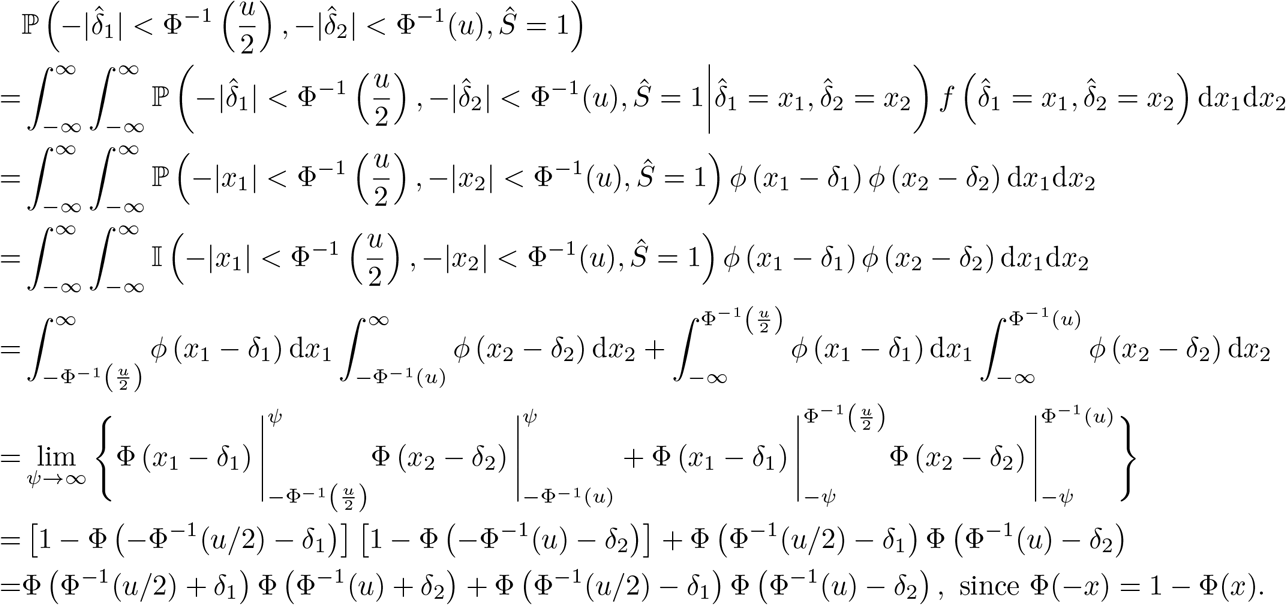

The fourth line holds because (*x*_1_, *x*_2_) are realizations of random variables, rather than being random themselves, so the probabilities evaluate to zero where their event is false and one where it is true. Following this example, we can evaluate Equation 3,

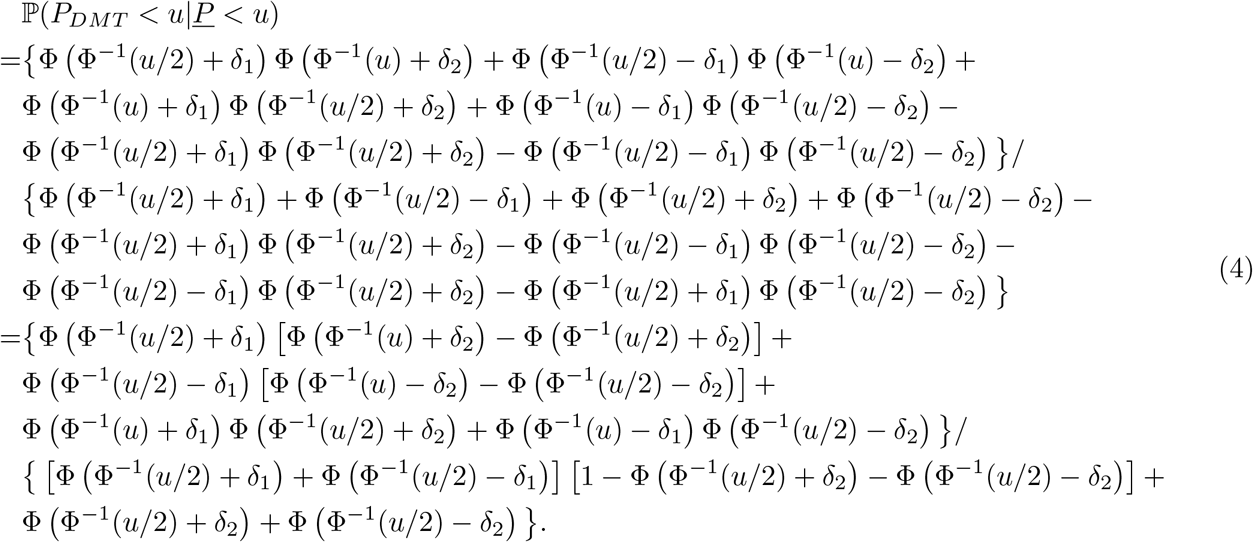

#### First partial derivative

To see where the maximum of Equation 4 may be for 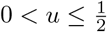, we first evaluate where

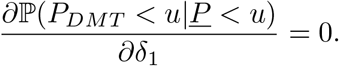

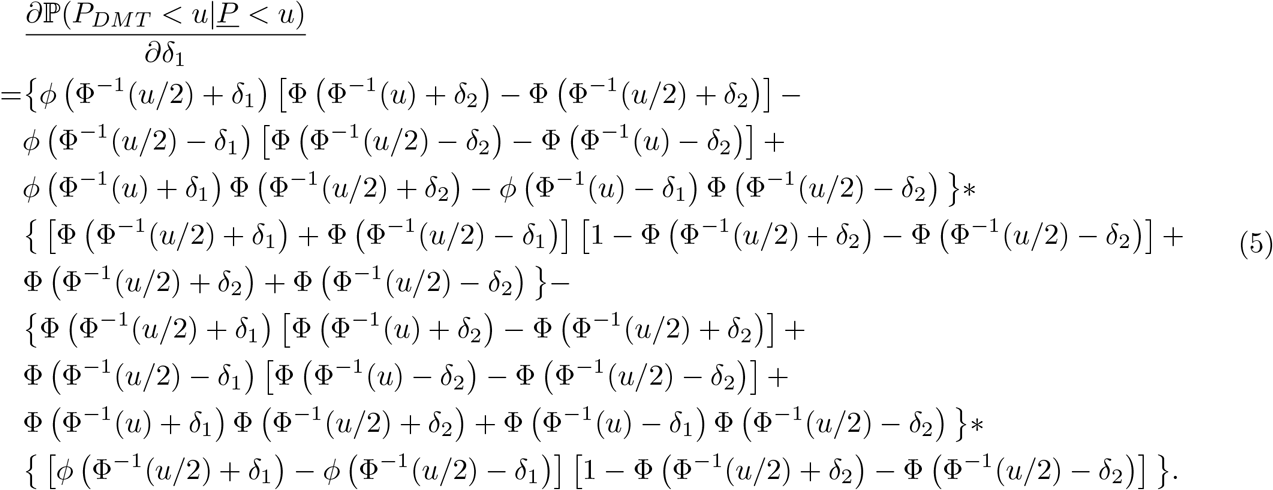

For fixed parameters off the boundary under 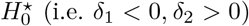, and 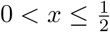, we have the following limits for *i* = 1, 2 as the sample size (and thus the magnitude of *δ*_*i*_) grows without bound,

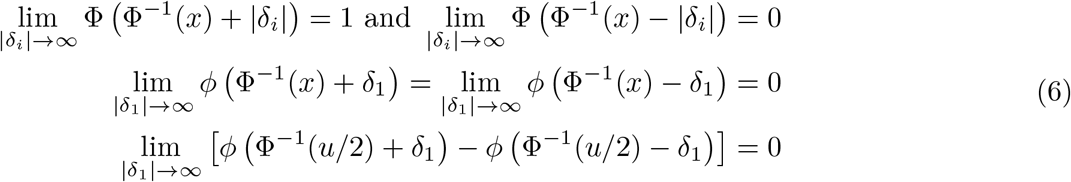

The second line in Equation 6 can be seen to hold based on the leading terms, which grow as exp 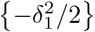.

The third line in Equation 6 is proportional to

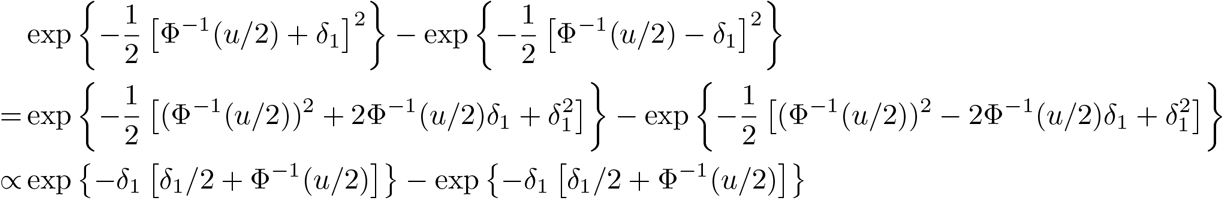

which is also dominated by exp 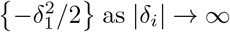 as *δ*_*i*_*→∞* or equivalently as *n→∞*. Given the limits in Equation 6, for parameters off the boundary under 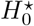 each term in Equation 5 goes to zero as the sample size increases by the product limit rule.

Thus off the boundary, 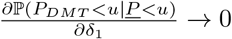 when *n*_1_ *→ ∞* and by symmetry of ℙ(*P*_*DMT*_ *< u*|*<u>P</u> < u*), also 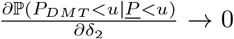. Thus, off the boundary both partial derivatives go to zero when (*n*_1_, *n*_2_) *→* (*∞, ∞*) and ℙ (*P*_*i*_ *< u*) *→* 1 for *i* = 1, 2 because of Equation 1, so ℙ(*<u>P</u> < u*) *→* 1. Whereas ℙ (*<u>P</u>< u*|*<u>P</u> <u*) *→* P(*Ŝ* = 1), and P(*Ŝ* = 1)*→*0 as (*n*_1_, *n*_2_) *→* (*∞,∞*), so ℙ(*P*_*DMT*_ *< u*|*<u>P</u> < u*)*→*0 and this is the global minimum. However, on the boundary, we have shown that ℙ(*P*_*DMT*_ *< u*|*<u>P</u> < u*)*→u* when only one parameter is zero as the sample size grows unbounded, which is the asymptotic maximum under 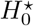, which completes the proof.

### Conditional p-value in finite sample sizes

We have the closed form expression of the conditional p-value ℙ (*P*_*DMT*_ *< u*|*<u>P</u> < u*) in Equation 4 for finite sample sizes, so we plot it below. We see that when *δ*_1_ is off the boundary in the left panels (here we show *δ*_1_ =*−*10) and *δ*_2_ *>* 0, conditional validity is easily satisfied, since the curve is far below the dashed line at *u*. When *δ*_1_ = 0 is on the boundary in the right panels and *δ*_2_ takes on some small values there is slight inflation but conditional validity holds when *δ*_2_ is near zero and when *δ*_2_ *>* 0 for large sample sizes, which is concordant with our mathematical proof.

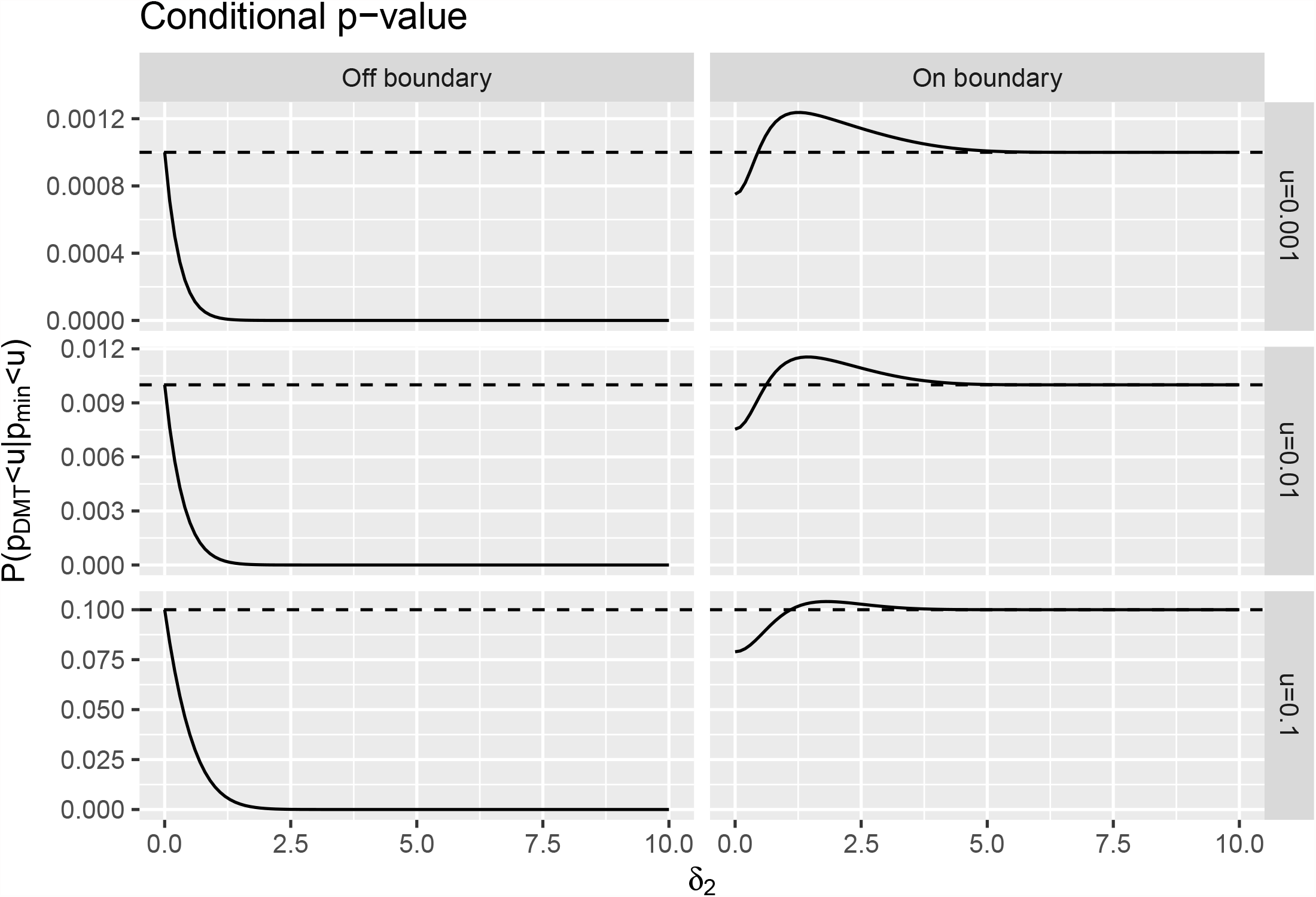

